# Early but not late exercise training in mice exacerbates hepatic inflammation in early NAFLD

**DOI:** 10.1101/2022.11.28.518192

**Authors:** Artemiy Kovynev, Zhixiong Ying, Joost Lambooij, Bruno Guigas, Patrick C.N. Rensen, Milena Schönke

## Abstract

Exercise effectively prevents obesity-related disorders, but it is unclear whether the beneficial health effects of exercise are restricted to unique circadian windows. Therefore, we aimed to study whether timing of exercise training differentially modulates the development and progression of non-alcoholic fatty liver disease (NAFLD), a disease currently estimated to affect over two billion people worldwide. We endurance-trained high fat-high cholesterol-fed NAFLD-prone male APOE*3-Leiden.CETP mice five times per week for eight weeks either in the early (ZT13) or in the late (ZT22) active phase and assessed the NAFLD score (histology) and hepatic inflammation compared to sedentary mice. Exercise training prevented an increase in body fat mass and fasting plasma glucose as expected, but neither early nor late training affected liver triglyceride or cholesterol content compared to sedentary mice, likely due to a very early stage of hepatic steatosis. In line, hepatic expression of de novo lipogenesis genes (e.g., Fasn, Srebp1c) was similarly downregulated by early and late training. However, exercise had a distinct time-dependent effect on hepatic inflammation, as only early training promoted an influx of pro-inflammtory cells into the liver paired with increased expression of the pro-inflammatory cytokines (e.g. Tnfa, Il1b). This data suggests that the timing of exercise is a critical factor for the effect on cardiometabolic disease development.

---

Two billion people worldwide are estimated to have non-alcoholic fatty liver disease (NAFLD), defined by excess hepatic fat. Commonly, the disease progression into non-alcoholic steatohepatitis (NASH) is characterized by the onset of liver inflammation following worsening steatosis [1]. This suggests that a reduction of liver inflammation – e.g. through lifestyle interventions such as exercise training – may only come secondary to a reduction of advanced steatosis. However, both the metabolic and inflammatory processes involved in NAFLD development are under circadian control and could hence respond differently to exercise at different times of day [2]. To investigate the time-of-day dependent effect of exercise training on NAFLD amelioration in the early disease stages we trained high-fat high-cholesterol (HFHC)-fed APOE*3-Leiden.CETP mice early or late in their active period. For this, animals were trained on a treadmill (17 m/min) for 1 hour at either *Zeitgeber* time (ZT)13 (E-RUN) or ZT22 (L-RUN) five days per week. Corresponding sedentary animals (E-SED and L-SED) were put into empty cages without bedding at the same time to control for the experienced stress. This mouse model was chosen due to its humanized lipid metabolism and its ability to develop all hallmarks of human NAFLD upon HFHC feeding [3].

Following 8 weeks of training, all mice had a similar body weight (Fig. 1A) and lean body mass (Fig. S1A), but both exercising groups had gained less fat mass than their sedentary counterparts, which only reached statistical significance in the comparison of the early groups (Fig. 1B), indicating a measurable exercise effect. Fasting plasma glucose levels, independently positively associated with the risk to develop NAFLD [4], were unchanged among the groups (Fig. 1C). Simultaneously, no differences in hepatic steatosis, NAFLD activity score and liver weight were observed between any of the groups (Fig. 1D-F), likely due to an overall limited treatment potential of early steatosis. Accordingly, liver lipid levels (total cholesterol, triglycerides and phospholipids) were unchanged between the exercising and sedentary groups regardless of the time of training (Fig. 1G-I), and so were the levels of plasma triglycerides and cholesterol (Fig. S1B-C).

**Figure 1.**
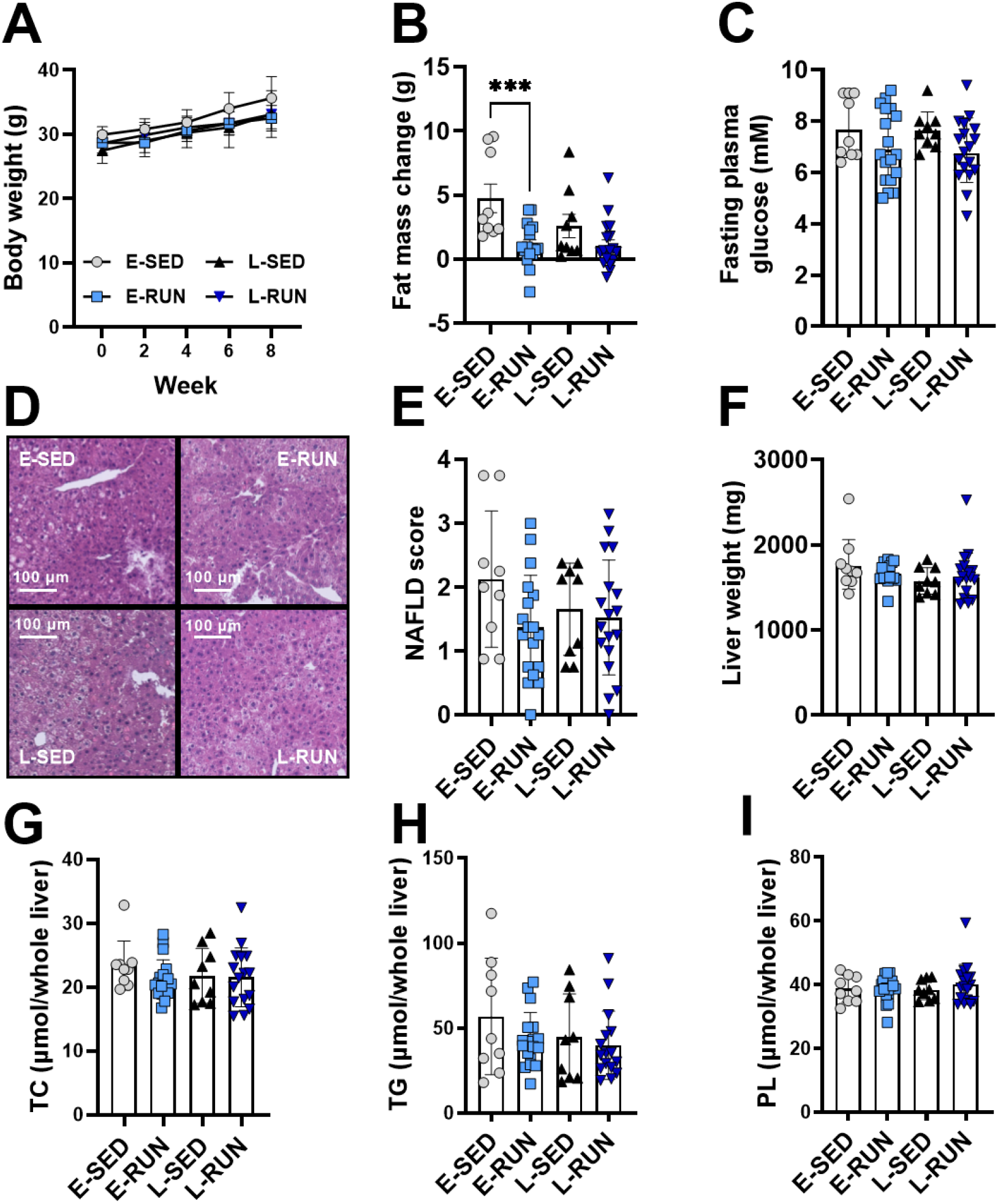
Early and late exercise training both improve body composition but do not alter early liver steatosis in APOE*3-Leiden.CETP mice. Over 8 weeks of treadmill training, body weight (A) and changes of fat mass (B) were monitored and fasting plasma glucose was measured after 8 weeks (C). Representative images of H&E-stained liver sections are shown (D) that were used to assess the NAFLD score (E). Liver weight (F), total liver cholesterol (G), triglyceride (H) and phospholipid (PL) content (I) were assessed after 8 weeks. ***P<0.001 in one-way ANOVA, n=9-18.

Surprisingly, however, exercise training had a time-of-day specific impact on liver inflammation, challenging the notion that hepatic inflammation can only be modulated later on in the disease through the reduction of steatosis. In livers collected at the same circadian timepoint (4 hours into the dark phase at ZT16; 17 and 26 hours after the last exercise bout for L-RUN and E-RUN, respectively), flow cytometry of the hepatic immune cells revealed a significant and unexpected increase in the number of leukocytes, neutrophils and monocytes with early exercise training (Fig. 2A-C). Late exercise, on the other hand, had no effect on liver immune cell populations. The increase of these cell populations particularly with early exercise may signify disease acceleration as infiltrating neutrophils are associated with early NAFLD development and progression into NASH [5, 6]. In line, infiltrating monocytes, that are recruited to the liver through hepatocyte-derived stress signals (such as IL-1 β and TNFα), promote NASH development once they differentiate into lipid-associated macrophages [7]. Interestingly, early exercise training also increased the number of natural killer (NK) cells in the liver (Fig. 2D). The contribution of these cells that are specialized on killing infected and tumor cells to NAFLD development and progression to NASH remains controversial, but they produce large quantities of pro-inflammatory cytokines such as IFNγ [8]. Taken together, early training leads to an inflammatory response in the liver characterized by an increase of pro-inflammatory and lipid damage-related cell populations.

**Figure 2.**
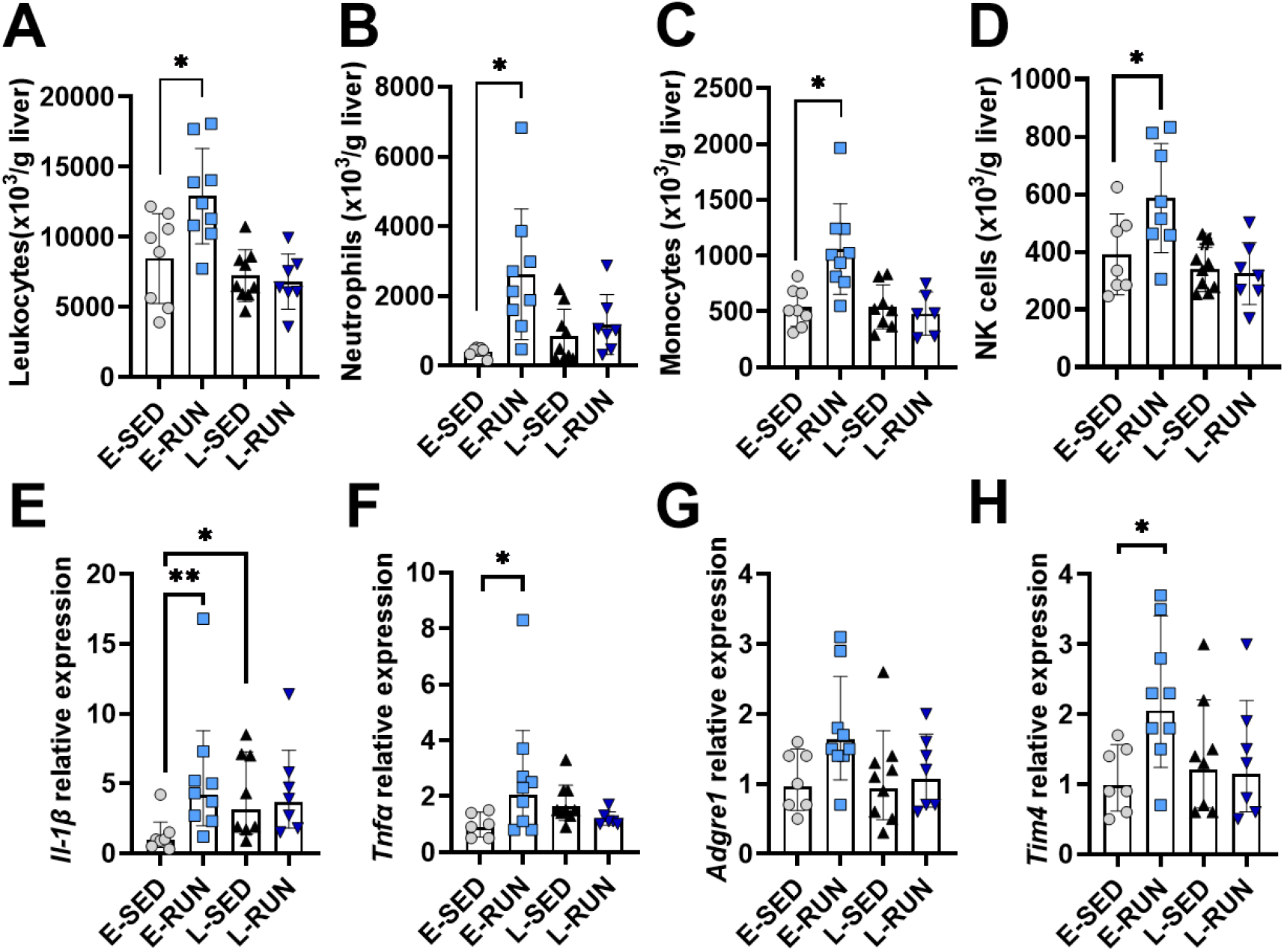
Early exercise promotes distinct changes to liver immune cell populations and inflammatory markers in early steatosis. The number of liver leukocytes (A), neutrophils (B), monocytes (C) and NK cells (D) was determined after 8 weeks of treadmill training using flow cytometry. The gene expression of *Il-1β* (E), *Tnfα* (F), *Adgre1* (G) and *Tim4* (H) was assessed in the isolated liver immune cells and shown relative to the expression levels of E-SED. *P<0.05, **P<0.01 in one-way ANOVA, n=7-9.

An overall increase in liver inflammation with early exercise training was also confirmed by gene expression analyses in the isolated liver immune cells via quantitative polymerase chain reaction (qRT-PCR). The expression of the pro-inflammatory markers *Tnfα* and *Il-1β* and *Tnfα* was increased only with early but not with late exercise training (Fig. 2E-F). Similarly, the expression of the macrophage marker *Adgre1* (F4/80) tended to be increased and *Tim4,* a marker of monocyte-derived Kupffer cells [9], was increased only with early training (Fig. 2G-H). In line, in whole liver tissue early exercise training also increased the expression of *Tnfα, Il-1β* and *Adgre1* (Fig. S1D-F).

From these findings, it is unclear whether the observed increase of liver inflammation in early NAFLD with early exercise training is beneficial or detrimental. Possibly, by stimulating liver inflammation, early exercise training activates an earlier immune response that contributes to disease resolution. On the other hand, it has been shown that early exercise can acutely worsen metabolic diseases as seen in people with obesity and type 2 diabetes where early high intensity cycling elicited unfavorable blood glucose spikes that did not occur with late exercise [10]. Accordingly, our findings could indicate that early exercise training accelerates disease progression while late exercise potentially only affects liver steatosis and inflammation at a later disease stage. However, while not affecting liver lipid levels, the hepatic gene expression of *Srebp1c,* the mediator of insulin-induced fatty acid synthesis, was similary downregulated with early and late training (Fig. S1G), suggesting that the regulation of metabolic and inflammatory disease drivers may not be synchronized. Future studies need to investigate how this immuno-modulatory exercise effect in early steatosis translates to advanced disease stages and to human NAFLD and NASH. Notably, while it is believed that the initiation of inflammation happens after the worsening of steatosis, we observe distinct inflammatory modulation already at an early stage of the disease with a low NAFLD score and low grade steatosis. This may present a previously underappreciated inflammation-targeted treatment opportunity in a large part of the population at risk for NASH.

In summary, we demonstrate that early and late exercise training in a mouse model of NAFLD differently influenced liver inflammation in early steatosis. While both early and late exercise prevented a gain of body fat mass in comparison to sedentary animals, an unexpected increase in liver inflammation was observed with early exercise training.

## Acknowledgements

We thank Trea Streefland and Reshma Lalai (Div. of Endocrinology, Dept. of Medicine, LUMC, Leiden, the Netherlands) for their excellent technical assistance. We furthermore thank Lars Hoeve, Sjahnaaz Bholai and Jack Brouwer for their technical contributions. This study was financed by a grant from the Novo Nordisk Foundation to M.S. (NNF18OC0032394). Z.Y. was supported by a full-time PhD scholarship from the China Scholarship Council.

