## Supplementary methods and figure for "Early but not late exercise training in mice exacerbates hepatic inflammation in early NAFLD"

### Animal study

NAFLD-prone male APOE*3-Leiden.CETP mice were obtained as previously described [1, 2]. At 8-14 weeks of age, mice were group-housed in light-tight cabinets at 21°C under 12/12-hour light-dark conditions with cage enrichment. The cabinets were illuminated with white fluorescent light (intensity: 200-250 lux). All training bouts and experiments took place under dim red light illumination during the active period of the mice. Mice were fed a NAFLD-inducing high fat high cholesterol (HFHC) diet containing 60% fat and 1% cholesterol (Ssniff, Soest, Germany) *ad libitum*. After a dietary habituation period of 1 week, animals were block-randomized into three groups based on body weight, fat mass, lean mass (assessed by EchoMRI 100-Analyzer; EchoMRI, Houston, Texas), plasma triglycerides (TG), and total plasma cholesterol (TC). Group sizes were based on previous studies with this animal model. Animals that did not respond to the HFHC feeding with increased plasma lipids were excluded. Early dark phase running ("Early runners", E-RUN, n=18) took place one hour after lights off at *Zeitgeber* time (ZT) 13-14, late dark phase running ("Late runners", L-RUN, n=18) one hour before lights on at ZT 22-23 and sedentary mice (SED, n=18) did not train. The sedentary control group was split up, half (n=9) was housed in the same light cabinet as E-RUN and half (n=9) in the same light cabinet as L-RUN and they are shown as E-SED and L-SED. Body weight of all mice was assessed weekly and body composition, unfasted TG and TC again at the end of the study. All animals were killed at ZT17 17-26 hours after the last exercise bout via $CO_2$ inhalation and perfused with ice-cold PBS for 5 minutes before tissues were collected. All animal experiments were performed in accordance with the Institute for Laboratory Animal Research Guide for the Care and Use of Laboratory Animals and were approved by the National Committee for Animal experiments by the Ethics Committee on Animal Care and Experimentation of the Leiden University Medical Center.

Exercise training

Mice were trained on a rat treadmill with five lanes (MazeEngineers, Skokie, Illinois), allowing 3-4 cage mates to run on one lane together which was observed to improve the motivation of the mice to run. No electric shocks were used; instead mice were gently nudged back on the belt with a brush when they stopped running. After three days of treadmill acclimatization with increasing speed and running duration the mice were trained five times per week for one hour (15 min warm-up at 6-15 m/min, 15 min at 15 m/min and 30 min at 17 m/min; in total 899 m per bout for each mouse) for a total of eight weeks. To account for handling stress and the unknown environment that the training mice experienced, all SED mice were moved in groups into empty cages without bedding standing close to the treadmill for the duration of the running bout.

Plasma and liver lipid measurements

Plasma TG and TC were measured after the dietary habituation period directly before the start of the training and after 8 weeks of training. Liver lipids were extracted from snap-frozen liver tissue using the protocol from Bligh and Dyer [3]. TG and TC concentrations where measured using the Total Triglyceride and Total Cholesterol assay kits (both Roche Diagnostics, Almere, The Netherlands) as described previously [4], and PL was measured using the Phospholipid Reagent kit (Instruchemie, Delfzijl, The Netherlands).

Liver immune cell isolation and flow cytometry

Liver tissue was collected in ice-cold RPMI 1640+Glutamax (Thermo Fisher Scientific, Waltham, MA, USA), minced and digested for 45 min at 37°C using collagenase type IV from *Clostridium histolyticum* (1 mg/mL in RPMI 1640+Glutamax; Sigma-Aldrich, St. Louis, MO, USA), 2000 U/mL DNase (Sigma-Aldrich, St. Louis, MO, USA) and 1 mM $CaCl_2$ as previously described [5]. The digested tissues were passed through 100 µm cell strainers and washed with PBS supplemented with 0.5% BSA and 2 mM EDTA (PBS/BSA/EDTA). The samples were spun down (530 x g, 10 min at 4°C) and the pellet resuspended in

PBS/BSA/EDTA and centrifuged again at 50 x g to pellet the hepatocytes (3 min at 4°C). The leukocyte-containing supernatant was collected and spun down (530 x g, 10 min at 4°C) before the cell pellet was treated with erythrocyte lysis buffer (0.15 M $NH_4Cl$; 1 mM $KHCO_3$; 0.1 mM $Na_2EDTA$) for 2 min at room temperature. After washing with PBS/BSA/EDTA, the leukocytes were isolated using magnetic-activated cell sorting (MACS) using LS columns and CD45 MicroBeads (35 µL beads per liver; Miltenyi Biotec, Bergisch Gladbach, Germany) according to the manufacturer's protocol. Isolated CD45+ cells were counted and stained with Zombie NIR (Biolegend, San Diego, CA, USA) for 20 min at room temperature followed by fixation with 1.9% paraformaldehyde (Sigma-Aldrich, St. Louis, MO, USA) for 15 min at room temperature after which the fixed leukocytes were further processed for flow cytometry. For this, the isolated CD45+ cells were incubated with a cocktail of antibodies against XCR1, CD11c, CD19, Ly6G, F4/80, MHC-II, CD45, CLEC2, Siglec-F, CD64, CD8, NK1.1, CD11b, CD4, CD90.2, Ly6C, CD3, CD36, CD9 and TIM4 in PBS/BSA/EDTA supplemented with True-Stain monocyte blocker (Biolegend, San Diego, CA, USA) and Brilliant Stain Buffer Plus (BD Biosciences, Franklin Lakes, NJ, USA) for 30 min at 4°C. The stained samples were measured by spectral flow cytometry using a Cytek Aurora spectral flow cytometer (Cytek Biosciences, Fremont, CA, USA). Spectral unmixing of the flow cytometry data was performed using SpectroFlo v3.0 (Cytek Biosciences, Fremont, CA, USA). Gating of flow cytometry data was performed using FlowJoTM v10.8 Software (BD Biosciences, Franklin Lakes, NJ, USA). Dimensionality reduction by means of Uniform Manifold Approximation and Projection (UMAP) was performed using OMIQ data analysis software (Omiq inc, Santa Clara, CA, USA).

<u>Histological analyses</u>

Fresh liver tissue was fixed in 4% paraformaldehyde, embedded in paraffin and sectioned into 5 µm thick sections for H&E staining. The NAFLD activity score was determined following a clinically utilized scoring system [6] that ranges from 0-7, and is evaluated semi-quantitatively through three criterions: steatosis (0-3), lobular inflammation (0-2), and hepatocellular ballooning (0-2).

<u>Gene expression analysis</u>

RNA was isolated from the above isolated immune cells that had been frozen following isolation as well as freshly frozen liver tissue using TRIzol RNA isolation reagent (Thermo Fisher, Waltham, Massachusetts). Following reverse transcription with M-MLV Reverse Transcriptase (Promega, Madison, Wisconsin), qRT-PCR was performed using SYBR Green (Promega). The primers used were:

| Gene | Forward primer (5' - 3') | Reverse primer (5' - 3) |
|---|---|---|
| *Adgre1* | CTTTGGCTATGGGCTTCCAGTC | GCAAGGAGGACAGAGTTTATCGTG |
| *Il1b* | GCAACTGTTCCTGAACTCAACT | ATCTTTTGGGGTCCGTCAACT |
| *Rplp0* | GGACCCGAGAAGACCTCCTT | GCACATCACTCAGAATTTCAATGG |
| *Srebp1c* | AGCCGTGGTGAGAAGCGCAC | ACACCAGGTCCTTCAGTGATTTGCT |
| *Tim4* | AGAGACACAAGAGGCCAGACA | TACAACCAGACAGGACCCACA |
| *Tnfα* | AGCCCACGTCGTAGCAAACCAC | TCGGGGCAGCCTTGTCCCTT |

Expression of genes of interest was normalized to expression of the housekeeping gene *Rplp0* and is shown relative to E-SED.

<u>Statistical analyses</u>

All individual data points are shown or data are otherwise expressed as mean ± SEM. Statistical analyses were performed using GraphPad Prism 9.01 (GraphPad, La Jolla, California) and one-way or two-way ANOVAs followed by Tukey's multiple comparisons test were used where appropriate. In case of missing values, mixed-effects analysis instead of two-way ANOVA was performed. Statistical outliers were removed after identification by Grubb’s test. Differences between groups were considered statistically significant if P<0.05 (*), P<0.01 (**), P<0.001 (***) or P<0.0001 (****).

**Supplementary Figure 1. Lean mass changes, plasma lipids, plasma glucose levels and liver gene expression after 8 weeks of early or late exercise training.** Body lean mass was measured before and after 8 weeks of exercise training (A). Plasma triglycerides (B), total plasma cholesterol (C) and liver gene expression of *Tnfα* (D), *Il-1β* (E), *Adgre1* (F) and *Srebp1c* (G) were assessed after 8 weeks. Gene expression is shown relative to E-SED. *P<0.05, **P<0.01 in one-way ANOVA, n=9-18.

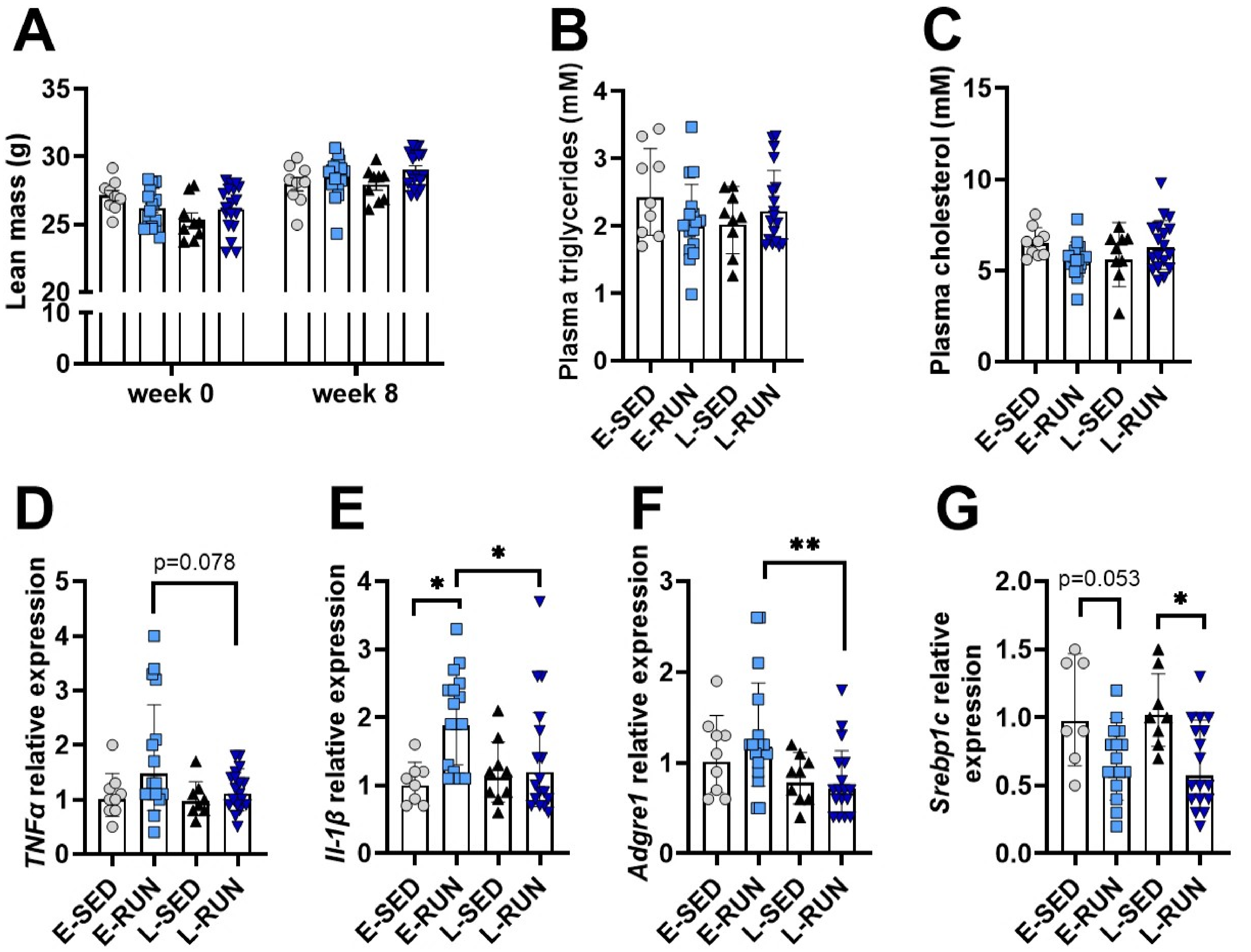

A
Lean mass (g)
35
30
25
20
10
0
week 0
week 8
B
Plasma triglycerides (mM)
4
3
2
1
0
E-SED
E-RUN
L-SED
L-RUN
C
Plasma cholesterol (mM)
15
10
5
0
E-SED
E-RUN
L-SED
L-RUN
D
TNFα relative expression
p=0.078
5
4
3
2
1
0
E-SED
E-RUN
L-SED
L-RUN
E
Il-1β relative expression
*
*
4
3
2
1
0
E-SED
E-RUN
L-SED
L-RUN
F
Adgre1 relative expression
**
3
2
1
0
E-SED
E-RUN
L-SED
L-RUN
G
Srebp1c relative expression
p=0.053
*
2.0
1.5
1.0
0.5
0.0
E-SED
E-RUN
L-SED
L-RUN
